# Effects of temperature on development, reproduction and size of *Trichogramma achaeae*: implications for biological control

**DOI:** 10.1101/2020.12.14.422618

**Authors:** Long Chen, Jesper Givskov Sørensen, Annie Enkegaard

## Abstract

The performance of biological control agents (BCAs) in outdoor crops is strongly regulated by ambient temperature. Understanding the thermal biology of BCAs and manipulating their thermal performance could improve biological control efficacy. In this study, the effects of temperature on several life history parameters (longevity, fecundity, development time, wing size) of the recently commercialised egg parasitoid *Trichogramma achaeae* Nagaraja & Nagarkatti (Hymenoptera: Trichogrammatidae) was examined. First, parasitoids were reared at 23 °C and tested in the laboratory at four constant temperatures (15, 20, 25 and 30 °C). Results demonstrated that temperature significantly altered all above parameters. Second, developmental acclimation was applied to manipulate the laboratory performance. Parasitoids were allowed to develop at either of the above four temperatures and their performance were compared at 23 °C. Results showed that developmental acclimation had a significant impact on fecundity, development time and wing size but not on female longevity. Our results have implications for improving the performance of *T. achaeae* in mass production and for its application for biological control under different thermal conditions.

**Highlights:**

- The temperature dependent performance of *Trichogramma achaeae* was characterised
- Acclimation significantly influenced fecundity, development and body size
- The overall performance was not improved by acclimation
- The female fecundity could be a proxy for the overall performance of *T. achaeae*

## 1. Introduction

Increasing concerns of potential hazards to the environment and human health from pesticide use has promoted development of biological control during the past decades (van Lenteren, 2012; van Lenteren et al., 2018). For augmentative biological control, which rely on releases of mass produced natural enemies, the quality of the released individuals are of utmost importance, with quality ultimately being defined as the ability to achieve a successful control under field conditions (van Lenteren, 2003). Quality is often approximated by laboratory measures of one or several life history parameters, such as fecundity, sex ratio, longevity, body size or host-searching ability in addition to tolerance to stresses under conditions in mass production (Sørensen et al., 2012; van Lenteren et al., 2003). It is assumed (but rarely verified) that organisms with good performance in one or several of these proxy parameters also perform better under field conditions.

The assumption of quality of biological control agents based on simple laboratory parameters can be questioned for two main reasons. Firstly, the control efficacy relies on the performance of living organisms, which is influenced by the ambient biotic and abiotic environment. Most of the organisms used in biological control are ectothermic insects, and their physiology and behaviour strongly rely on ambient temperature. In field application where temperature fluctuates over space and time, the control efficacy varies and might be strongly limited by unfavourable conditions (Pekár and Hubert, 2008; Tullett et al., 2004). Secondly, mass-rearing of insects could lead to undetected deteriorations in field quality. Mass-rearing of biological control agents (BCAs) usually occur at artificial environmental conditions with high constant temperature and, often, artificial diets to maximise generation turnover (Mackauer, 1976). Under these conditions, BCAs could experience potential laboratory adaptation, i.e. inadvertent selection for particular parameters (e.g. decreased development time or high fecundity) at the expense of other traits. If these are used as quality measures, such populations could be perceived as having high laboratory quality, while field performance might be very different (Gariepy et al., 2014). Furthermore, inbreeding and drift lead to decreased genetic variation and inbreeding depression (Sørensen et al., 2012), and would further contribute to quality-associated poor performance in the field (van Lenteren et al., 2003).

To improve quality of BCAs intended for application at specific target temperatures, one obvious way is to explore the thermal plasticity or acclimation. Thermal plasticity is the capacity of an organism to express multiple phenotypes in response to different temperatures thus allowing it to cope with stress by physiological and behavioural modifications (Angilletta 2009). Thus, insects with high thermal plasticity would be predicted to have better performance at fluctuating or unfavourable climatic conditions (Chevin and Hoffmann, 2017). Plastic modifications could potentially induce changes in quality related performance, for example, storing developing parasitoids at low temperatures may result in reduced reproduction and longevity (Colinet and Boivin, 2011). Many studies have suggested that organisms could increase their performance in environments close to their developmental or acclimation temperatures, as compared to organisms acclimated at other temperatures (Wilson and Franklin, 2002; Angilletta, 2009). These studies implied the possibility to acclimate BCAs and further improve their performance at particular temperature regimes in the field.

Acclimation has been studied for several natural enemies and some studies have supported that acclimation may increase performance under certain conditions. For example, acclimation alter the survival of insects at extreme temperatures (Chanthy et al., 2012; Hughes et al., 2010). Research on the predatory mite *Gaeolaelaps aculeifer* Canestrini (Acari: Laelapidae) showed that cold acclimation increased cold and starvation tolerance (Jensen et al., 2017). Similarly, cold reared (15 °C) two-spotted ladybird (*Adalia bipunctata* Linnaeus (Coleoptera: Coccinellidae)) consumed more aphids at 15 °C than they did when reared at 20 or 25 °C (Sørensen et al., 2013). The two above studies also pointed to an associated cost of acclimation with reduced performance at opposite temperatures or reduced predation rate or reproduction at certain temperatures. However, acclimation could still benefit the overall performance of BCAs, for instance by allowing establishment of cold acclimated predatory mites at cold areas, particularly when costs are identified and quantified (Hart et al., 2002). For short-lived insects, such as the egg parasitoids *Trichogramma* spp., the short life span limits the scope for sensing and plastically responding to environmental change, as these have been shown to be time-consuming processes (DeWitt et al., 1998). Therefore, developmental acclimation could potentially improve their biological control efficacy in unfavourable environment.

*Trichogramma achaeae* Nagaraja & Nagarkatti (Hymenoptera: Trichogrammatidae) is a recently commercialised species mainly used to control tomato leaf miner *Tuta absoluta* Meyrick (Lepidoptera: Gelechiidae) in tomato (Cabello et al., 2012). Successful biological control requires the knowledge of biology and ecology of a potential agent (Bale et al., 2008). However, little information is available on the thermal performance of this species. In a previous study by Cascone et al. (2015), longevity and fecundity was investigated at three different temperatures, which is inadequate to fully understand the consequences of different temperatures for the performance of this species. In the present study, in order to understand the thermal biology of *T. achaeae*, we tested how different ambient temperatures alter quality-related parameters (longevity, fecundity, development time, wing size) of *T. achaeae*.

To further explore whether developmental acclimation can potentially improve the performance of *T. achaeae*, we tested how different developmental acclimation temperatures affected the above parameters. We hypothesised that (1) increasing ambient temperature would result in reduced longevity, development time and wing size as well as in increased fecundity up to an optimal temperature; (2) 25 °C would be optimal as reported for other *Trichogramma* species (Krechemer et al., 2015); and (3) parasitoids acclimated at their optimal temperature would achieve a higher performance (in terms of fecundity or similar) than parasitoids acclimated at other temperatures.

## 2. Materials and Methods

### 2.1 Experimental animals

*T. achaeae* was obtained from Bioline AgroSciences Ltd, UK. At delivery, *T. achaeae* was in the pupal stage inside host eggs (species not disclosed by the company) glued onto cardboard pieces. The shipped material was used directly without storage. Eggs of the Mediterranean flour moth (*Ephestia kuehniella* Zeller, Lepidoptera: Pyralidae) were used as host for *T. achaeae* in rearing and experiments. UV sterilised *E. kuehniella* eggs were received every second week in bulk in bottles from Koppert B.V., the Netherlands, and stored at 4 °C. Parasitoids were fed with 50% (v: v) organic honey solution as suggested by Singhamuni et al. (2015).

### 2.2 Laboratory rearing

Parasitoids were reared continuously for a period of 4 months. Received *T. achaeae* cards (F_0_) were maintained in semi-transparent plastic bottles (approx. 300 ml) in a climate room at 23 ± 1 °C, with a photoperiod of 12: 12 (L: D). Bottles were positioned horizontally in a glass aquarium at the bottom of which was placed a petri dish (diameter: 9 cm) with saturated NaCl solution to maintain humidity in between RH 70 ± 10 %. Initially, each bottle contained 1 card with approximately 2,500 parasitised host eggs. Upon emergence, newly emerged parasitoids (< 24 h old) were provided with honey solution as food and with fresh *E. kuehniella* eggs glued on 2 cardboard pieces (1.3 * 6.3 cm/card, approximately 500 eggs/cm^2^) for oviposition for 24 h. Hereafter, the cards were transferred to new bottles and kept in the rearing aquarium as described above for adult emergence after which the rearing cycle was repeated. The first parasitoids to be used for experimentation (Experiment 1) came from the F_1_-generation (i.e. after a full generation of rearing in the laboratory) to avoid systematic errors arising from the rearing conditions at the producer (Good, 1993).

### 2.3 Performance at ambient temperature (Experiment 1)

The experiments were conducted in climate chambers at constant temperatures of 15, 20, 25, and 30 ± 1 °C, RH 70 ± 10 %, photoperiod 12 L: 12 D. Experiments were divided into 2 parts based on the measurements at different life stages. Longevity and fecundity were measured directly for females of F_1_ and development time and wing size (as a proxy for body size) were measured for the offspring of F_1_. Figure 1 shows the workflow of the experiments.

**Figure 1.**
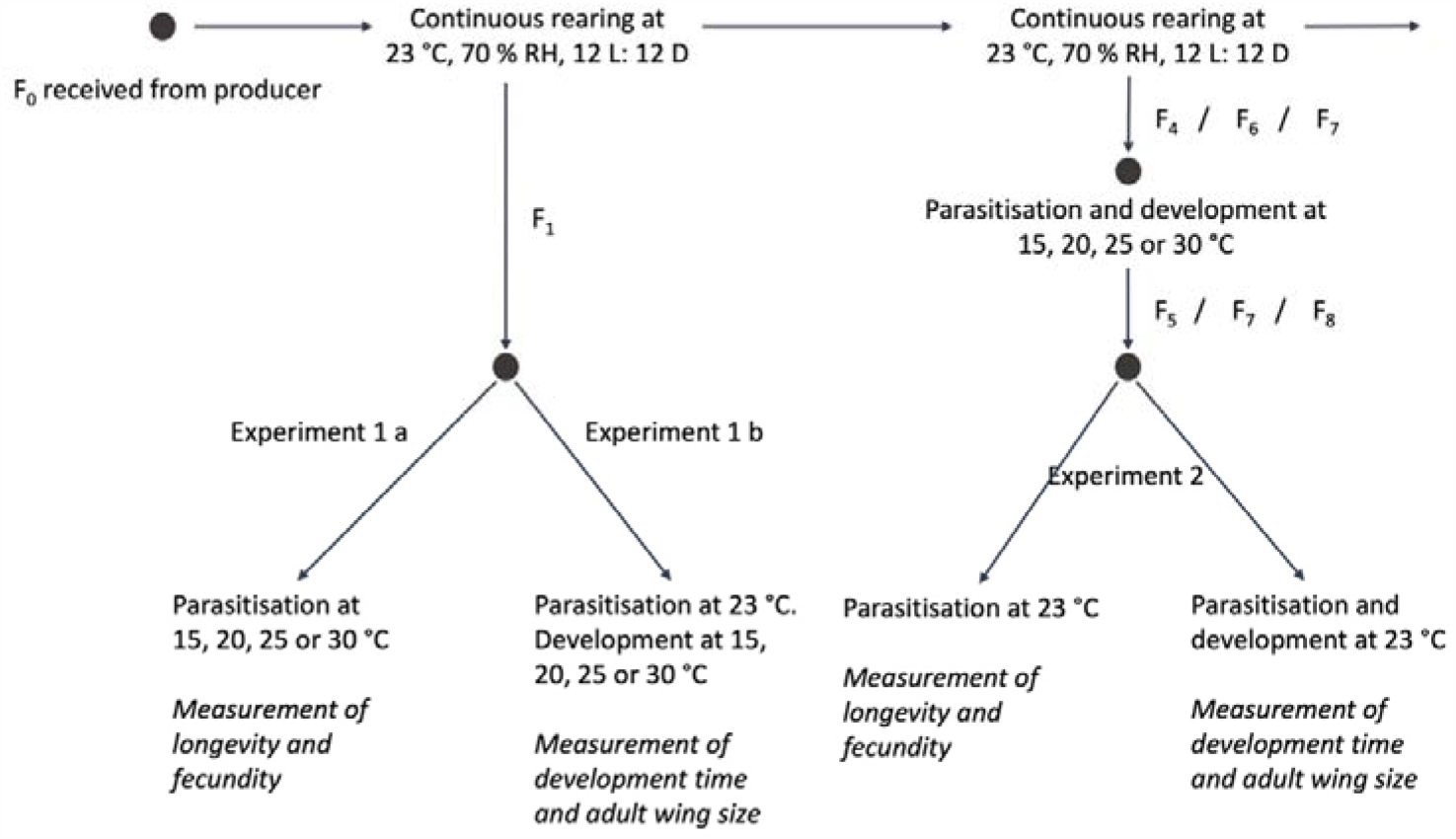
Experimental workflow. Parasitoids used in Experiment 1 originated from F_1_. Part of these parasitoids were directly measured for longevity and fecundity. The rest were given *E. kuehniella* egg for parasitisation and subsequent measurement of development time and wing size. Parasitoids used in Experiment 2 were taken from different generations (F_4_, F_6_, F_7_). Parasitoids were given *E. kuehniella* egg for parasitisation and subsequent development at either 15, 20, 25 or 30 °C. Emerged offspring were then measured at 23 °C for the same parameters as in Experiment 1

#### 2.3.1 Experimental 1a: longevity and fecundity

Prior to experimentation, egg cards with unhatched pupae were removed from the rearing bottles and emerged parasitoids were fed with honey and kept in bottles for additional 3 hours to ensure that all females were mated (Mills and Kuhlmann, 2000). For each treatment, 30 newly emerged females (< 24 h old) were randomly selected and individually maintained in 4 ml glass vials sealed with a piece of cotton. For each vial, an *E. kuehniella* egg card (0.5 * 1.3 cm, approximately 500 eggs/cm^2^) was set-in by tweezer and a fine drop of honey solution added to the bottom. Then all vials were horizontally placed in semi-transparent plastic boxes with a petri dish (diameter: 9 cm) of saturated NaCl solution to maintain humidity. Boxes were separately placed into 4 climate chambers at 15, 20, 25 and 30 ± 1 °C.

Longevity was recorded as the length of survival period of females. Recordings were made every 24 ± 2 h with individual longevities assigned to the mid-point between two observations. Daily recordings proceeded orderly from replicate 1 to 30 and from low temperature to high temperature to avoid systematic errors. Honey solution drops were checked daily and replenished at need.

Fecundity was measured as the number of new parasitoids emerging from *E. kuehniella* eggs that were parasitised during the first 5 days. At day 5, after the longevity recordings was done, *E. kuehniella* egg cards with these parasitised eggs were moved to new vials, and females were left in their original vials for continued measurement of longevity. Egg cards originating from the same temperature treatment were returned to this treatment for development of the parasitoids into adults. When emergence had ceased, all vials were placed in a freezer at - 24 °C to kill and store the parasitoids for later counting of the total number of emerged parasitoids.

#### 2.3.2 Experimental 1b: development time and wing size

Assessment of development time and wing size were made for offspring of F_1_-females at the four different temperature treatments. For each temperature, newly emerged females (< 24 h old) of F_1_ were offered 16 fresh *E. kuehniella* egg cards (0.5 * 1.3 cm) for 3 hours for parasitisation in a climate-controlled room at 23 ± 1 °C, RH 70 ± 10 % and 12: 12 L: D. The cards were subsequently randomly grouped into 4 replicates of 4 cards, transferred into vials, placed in similar boxes as in Experiment 1a and kept in climate chambers at the 4 different temperature treatments for parasitoid development. Egg cards were checked daily (every 24 ± 2 h) for emergence. When new parasitoids started to emerge, the egg cards were daily moved to new vials before the emerged parasitoids were killed and stored at -24 °C for later analysis. For each replicate, recordings ended when no emergence had been observed for 2 consecutive days. The development time was calculated as a weighted time according to the percentage of emergence at different dates. Sex-ratio was not measured as this is determined at the time of oviposition (Luck et al., 2001).

Wing size was measured for a total of 32 killed females per treatment (8 females per replicate). Females were randomly selected among the emerged parasitoids with the number collected from each emergence day being proportional to the daily emergence percentage out of the total. The length of the wings was measured from the node along with the connected vein to the edge (Figure 2). Measurements were made by taking photos of each female using a Leica microscope camera and determining wing length using ImageJ version 1.52 (Schneider et al., 2012).

**Figure 2.**
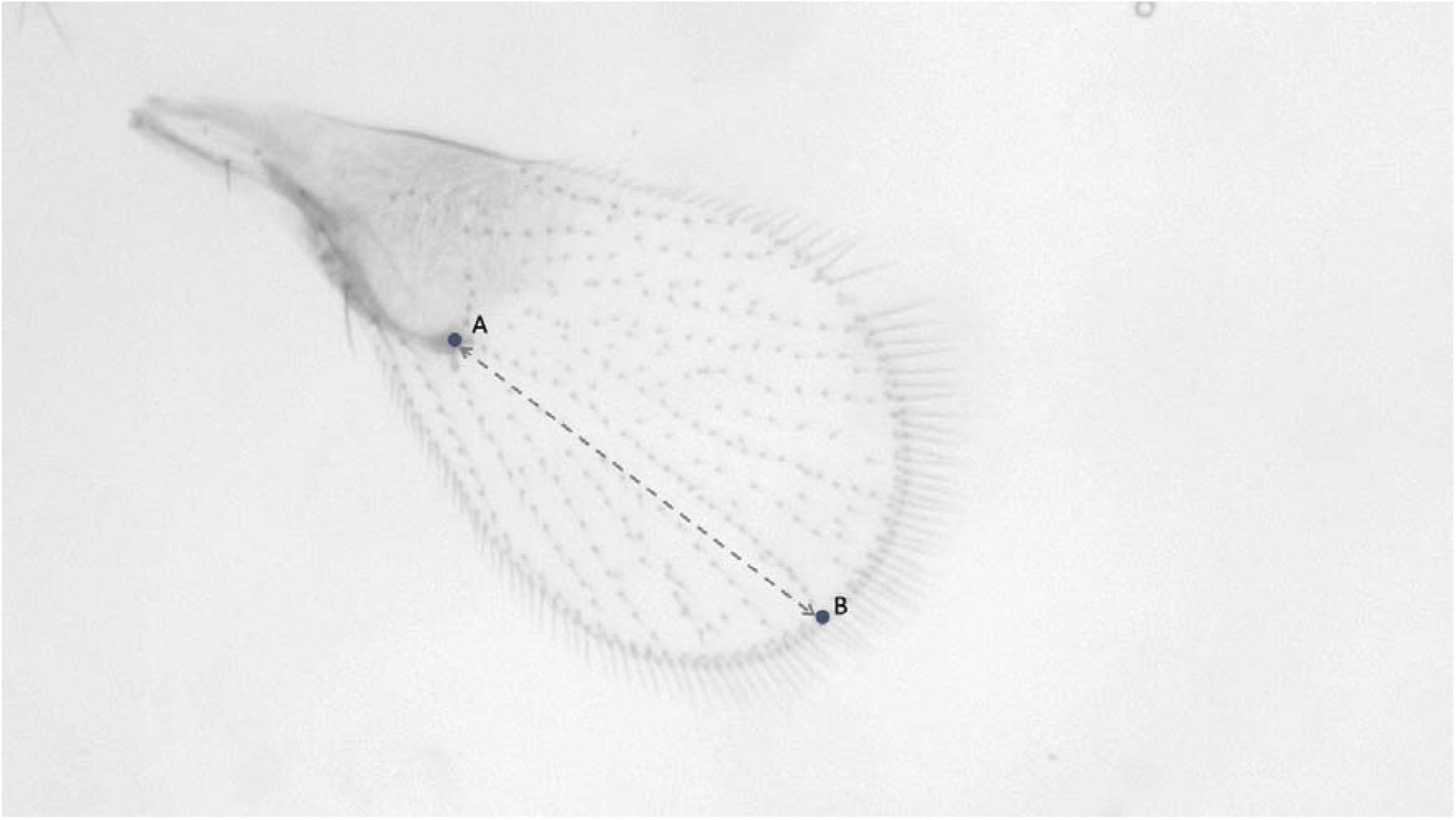
Wing size of *Trichogramma achaeae* was measured as the length of the dashed line connecting the middle point of the node (point A) to the conjunction (point B) of the edge of the forewing, i.e. approximating the length of the longest vein starting from the node.

### 2.4 Experiment 2: performance after developmental acclimation

To investigate the effects of developmental acclimation on longevity, fecundity, development time and wing size of *T. achaeae*, parasitoids were acclimated at four constant temperatures (15, 20, 25 and 30 ± 1 °C) for the whole developmental period, i.e. from parasitisation to emergence. Hereafter, parasitoids were tested at a constant temperature of 23 ± 1 °C (Figure 1). Parasitoids used for acclimation at the 4 different temperatures came from generation F_4_ (15 °C), F_6_ (20 °C) and F_7_ (25 and 30 °C) of the rearing (Figure 1).

For each treatment, newly emerged parasitoids (< 24 h old) were taken from the rearing and kept in semi-transparent plastic bottles (approx. 300 ml), fed with honey solution and provided with 2 fresh *E. kuehniella* egg cards (6.3 * 1.3 cm/card, 500 eggs/cm^2^). The bottles were placed in plastic box equipped as described for the rearing procedure (section 2.2.) and placed in a climate chamber at the respective acclimation temperature for parasitisation for 24 h. Hereafter, the eggs cards with the now parasitised *E. kuehniella* eggs were placed in 2 new bottles (one card/bottle) and kept at the same acclimation temperature for development until emergence. Upon the emergence of new parasitoids (< 24 h old), egg cards with unhatched pupae were removed and emerged parasitoids fed and kept in bottles at 23 °C for additionally 3 hours to ensure that all females were mated.

For each acclimation treatment, 30 females were randomly selected into vials (4 ml) to record longevity and fecundity at 23 °C. Another group of parasitoids was given 4 fresh *E. kuehniella* egg cards (0.5 * 1.3 cm) for 3 h for parasitisation at 23 °C. Then the parasitised eggs were maintained at 23 °C for measurement of development time and wing size. All recordings were made as described for Experiment 1 (section 2.3.1, 2.3.2). Due to low parasitisation of females acclimated at 15 °C, the sample size at this treatment was small. Thus, only a total of 27 females were used to measure wing size at the treatment of 15 °C compared to 32 females for the other temperature treatments. In addition, only 16 parasitoids per replicate were measured for development time at 15 °C, compared to more than 280 per replicate for the other temperature treatments.

### 2.5 Data analysis

All data analyses were carried out using R (Version 3.6.2) (R Core Team, 2019). Thermal effects on life history parameters were analysed using a general linear model (function lm). Firstly, all life history parameters were fitted with a simple linear model followed by Q-Q and residual plots to verify assumptions of normality of residuals and homogeneity of variances. Then second order poly-linear regression was performed when simple linear models did not fit the data well. Finally, one-way ANOVA analysis was used where parametric assumptions were met to explore the changes of temperatures on thermal performance of *T. achaeae*. Model selecting was determined by ANOVA test and Akaike information criterion (AIC) values. The lower temperature thresholds (T_0_) for development was estimated by linear regression of development rate on temperature as described by Davidson (1944).

## 3. Results

### 3.1 Performance at ambient temperature (Experiment 1)

Wing size and several life history characteristics of *T. achaeae* parasitising *E. kuehniella* eggs were significantly influenced by temperature. Thus, both longevity, development time and wing size decreased with increasing temperature, while fecundity increased with temperature (for a certain temperature range) (Table 1; Figure 3).

**Table 1.**
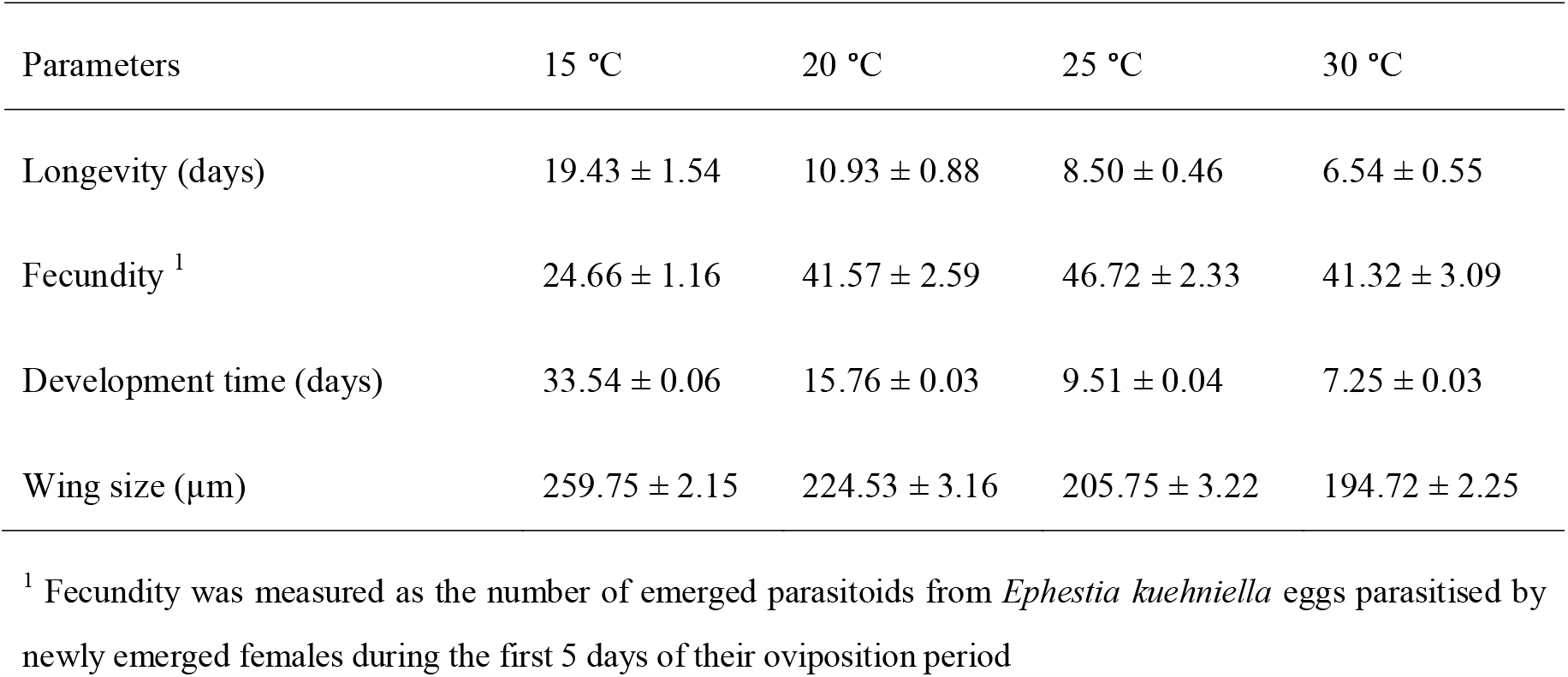
Life history parameters and wing size (mean ± SEM) of *Trichogramma achaeae* at 4 constant temperatures

**Figure 3.**
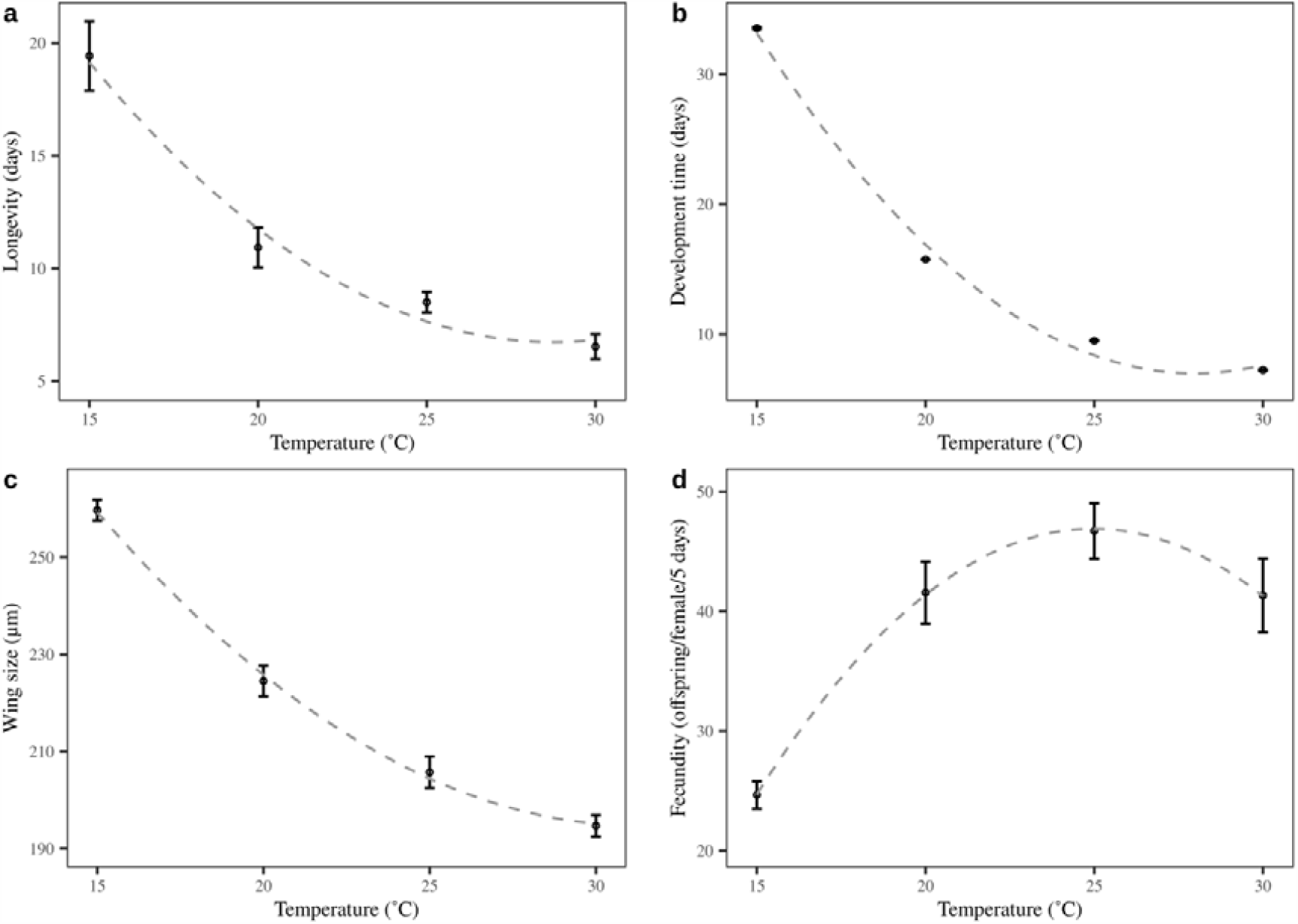
Means (± SEM) longevity (a), development time (b), wing size (c) and fecundity (d) of *Trichogramma achaeae* at four constant temperatures. Dashed lines are predicted values according to model 1

The temperature dependency for longevity, development time, wing size and fecundity could be described by a second order polynomial model:

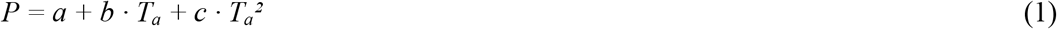

where *P* is parameter in question, *T*_*a*_ is the ambient temperature (°C) and *a, b* and *c* are constants.

For longevity, which varied between 7 and 19 days (Table 1), the model fitting (F_(2, 113)_ = 50.6, p < 0.001) gave the curve in Figure 3a and the following parameter estimates (± SEM): *a*: 61.039 ± 9.386, *b*: -3.779 ± 0.873, *c*: 0.066 ± 0.019.

The development of immature *T. achaeae* took from 7 to 34 days (Table 1) and the model fitting (F_(2, 13)_ = 960.5, p < 0.001) yielded the curve in Figure 3b and the following parameter estimates (± SEM): *a*: 128.545 ± 4.555, *b*: -8.687 ± 0.424, *c*: 0.155 ± 0.009. The lower temperature threshold (T_0_) for the juvenile development of *T. achaeae* was estimated as 11.0 °C (ANOVA, F_(3, 12)_ = 17290, p < 0.001).

For wing size, varying between 195 and 260 µm (Table 1), the following parameter estimates (± SEM) were obtained for the fitted model (F_(2, 125)_ = 162.7, p < 0.001; Figure 3c): *a*: 432.322 ± 26.591, *b*: -15.162 ± 2.473, *c*: 0.242 ± 0.055.

Finally, the fecundity of *T. achaeae*, ranging between 25 and 47 viable offspring per female for the first 5 days of the oviposition period (Table 1), was model fitted (F_(2, 113)_ = 24.4, p < 0.001) to give the curve in Figure 3d and the following parameter estimates (± SEM): *a*: - 92.278 ± 23.130, *b*: 11.148 ± 2.151, *c*: -0.223 ± 0.048.

### 3.2 Performance after developmental acclimation (Experiment 2)

Developmental acclimation had a significant influence on fecundity, development time and wing size of *T. achaeae* (Table 2). However, developmental acclimation did not affect female longevity (F_(3, 114)_ = 1.7, p = 0.17, mean ± SEM: 8.18 ± 0.33 days across all temperatures, Table 2).

**Table 2.**
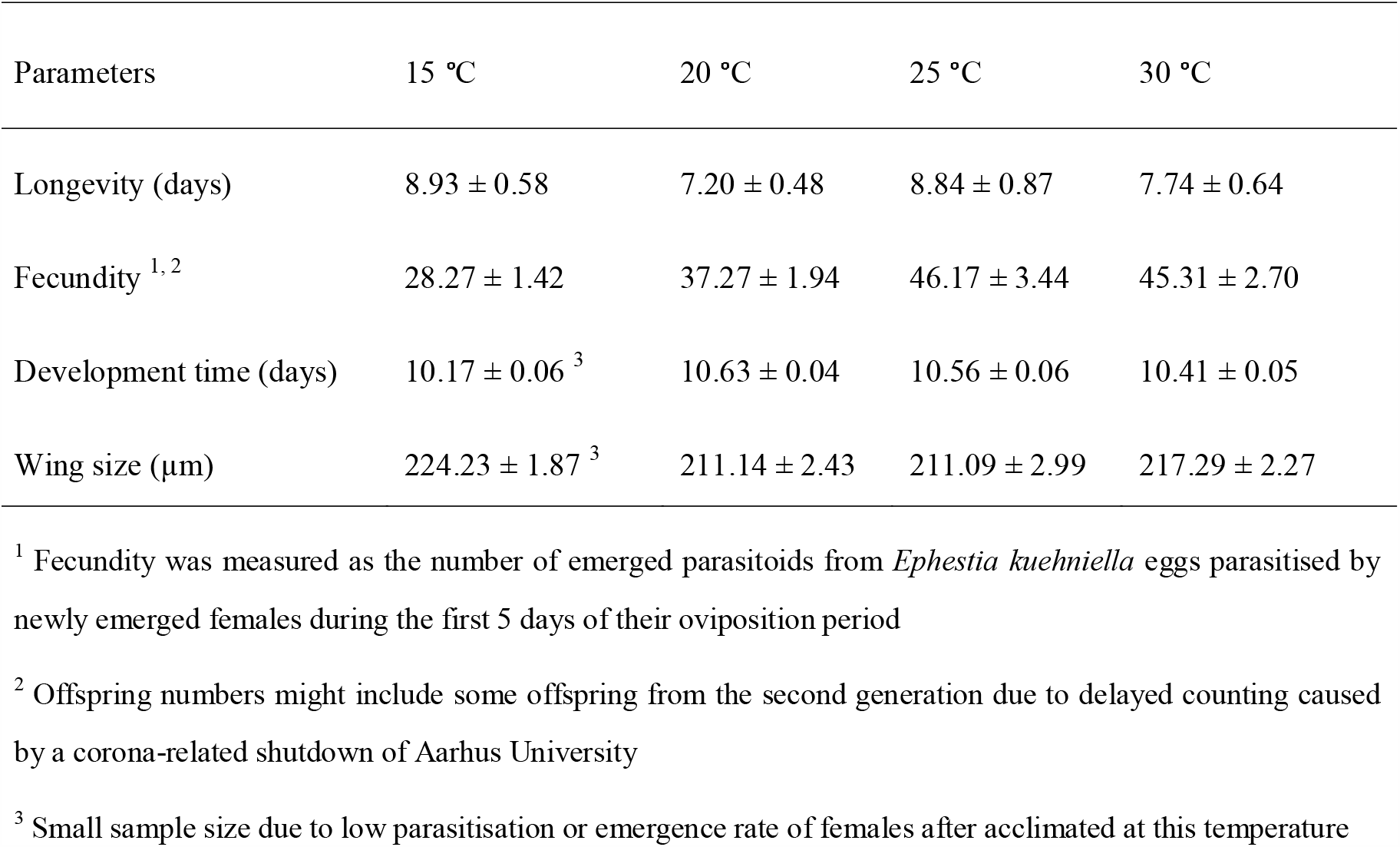
Life history parameters and wing size (mean ± SEM) at 23 °C for offspring of *Trichogramma achaeae* developmentally acclimated at 4 constant temperatures.

The influence of acclimation temperature on fecundity, development time and wing size could be described by a second order polynomial model:

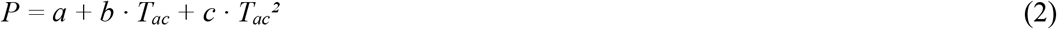

where *P* is parameter in question, *T*_*ac*_ is the acclimation temperature during development (°C) and *a, b* and *c* are constants.

The fecundity of *T. achaeae* expressed at 23 °C after developmental acclimation increased with acclimation temperature, varying between 28 and 46 viable offspring per female for the first 5 days of the oviposition period (Table 2). The model fit (F_(2, 115)_ = 16.8, p < 0.01) gave the curve in Figure 4a and the following parameter estimates (± SEM): *a*: -34.304 ± 23.989, *b*: 5.609 ± 2.233, *c*: -0.098 ± 0.049.

**Figure 4.**
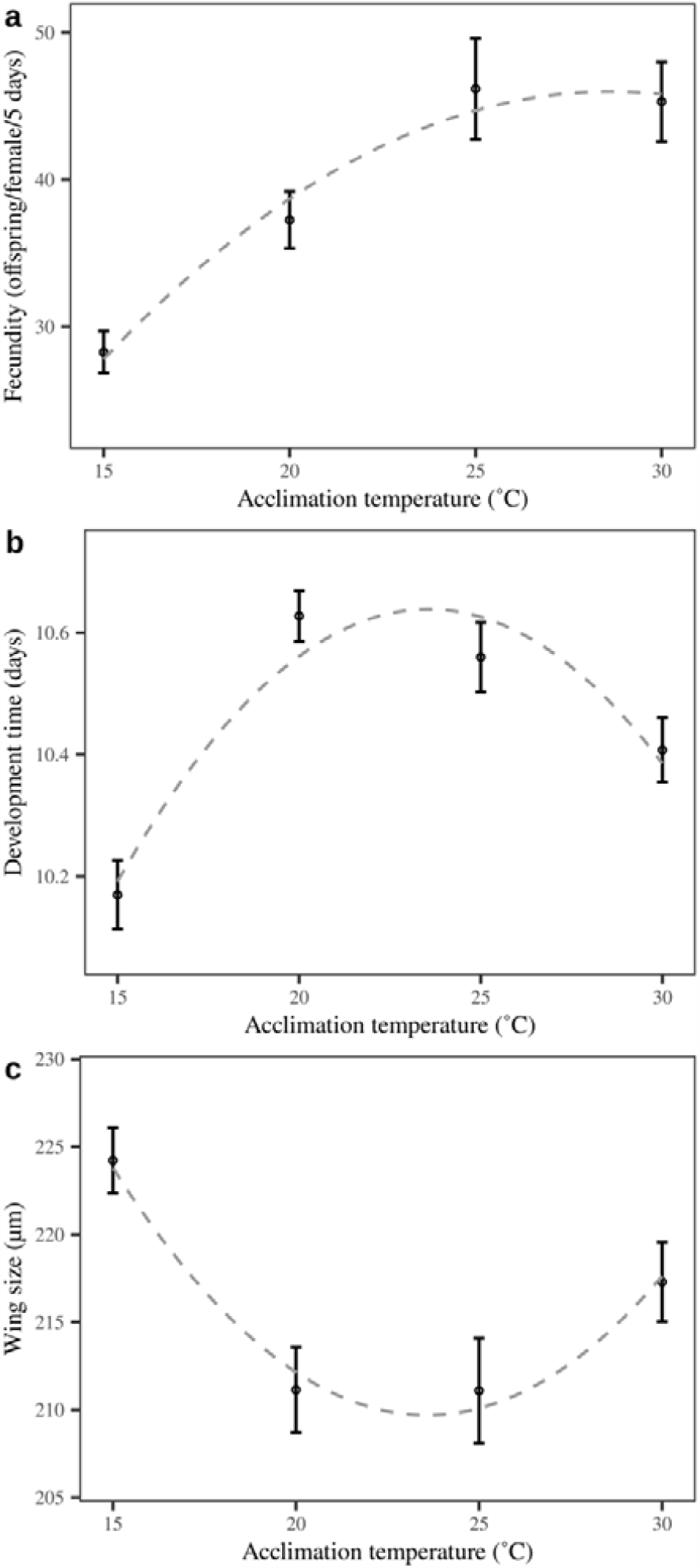
Means (± SEM) of three life history parameters, fecundity (a), development time (b) and wing size (c) of *Trichogramma achaeae* tested at 23 °C after developmental acclimation at four constant temperatures. Dashed lines are predicted values according to model 2

Acclimated immature *T. achaeae* had a development time at 23 °C varying between 10 and 11 days (Table 2). The model fit (Figure 4b) (F_(2, 13)_ = 17.0, p < 0.001) gave the following parameter estimates (± SEM): *a*: 7.254 ± 0.557, *b*: 0.287 ± 0.052, *c*: -0,006 ± 0.001.

Finally, the wing size of female offspring of *T. achaeae* that had undergone developmental acclimation ranged between 211 and 224 µm (Table 2). The model fit (F_(2, 118)_ = 8.5, p < 0.001) gave the curve in Figure 4c and the following parameter estimates (± SEM): *a*: 316.373 ± 24.828, *b*: -9.052 ± 2.278, *c*: 0.192 ± 0.050.

## 4. Discussion

Our characterisation of the temperature-dependency of the performance of *T. achaeae* is of relevance for mass production and field applications. Extended life span would increase the possibility for BCA establishment and maintenance of control efficacy for longer periods of time. Similarly, larger size is correlated with long life span, and high fecundity and body size directly affects field dispersal and searching ability (Kölliker-Ott et al., 2003; Beukeboom, 2018). In mass rearing, size and longevity could be optimised by keeping the rearing temperature low. However, at lower temperatures development time increase significantly which would present difficulties in maximising population turnover.

For parasitoids, fecundity directly represents the pest kill rate and links to field success (Coelho et al., 2016). In the present study, fecundity maintained a stable level at a broad temperature range (20 to 30 °C) making *T. achaeae* suitable for use in several geographical regions. However, a clear reduction of fecundity was observed when temperature decreased to 15 °C which is in line with observation for other *Trichogramma* species (Haile et al., 2002; Pizzol et al., 2010). The highest realised fecundity was achieved around 25 °C, which was similar to previous studies on other *Trichogramma* species (Krechemer et al., 2015). Overall, these findings partially supported our hypotheses 1) and 2).

Trade-offs were apparent between different temperature dependent characteristics of *T. achaeae*. Benefits from increased longevity and body size at low temperatures were thus offset by increased development time and reduced fecundity. Previous studies have reported that resource requirements for survival might leave less energy for reproduction (Hurd, 2001). Although body size generally has been found positively related to fecundity in insects (Honěk, 1993), this size-fecundity relationship is reversed across temperatures. Lower developmental temperature results in enlarged body size, however, lower temperatures are usually associated with reduced fecundity. The results of this study suggests that finding a temperature that balance trade-offs between different parameters is feasible. The data showed that fecundity was likely not constrained by a trade-off with longevity with temperature, as the life span period is almost sufficient for females to parasite and deplete their egg load (Özder and Kara, 2010; Scholler and Hassan, 2001). Therefore, compared to the indirect effects on biological control efficacy of development time and body size, the presented data point to fecundity as a general predictor of overall performance under different temperature regimes. Thus, we suggest that any temperature range in favour of fecundity is recommended for achieving high performance. Based on this study, temperatures around 25 °C would be a suitable temperature for *T. achaeae* to perform well.

Differing from most studies that have directly tested acclimation effects on insect performance at extreme temperatures, we tested acclimated *T. achaeae* at an intermediate temperature. Therefore, results in this study did not provide information as to whether acclimation might improve thermal tolerance, but has demonstrated that acclimation can alter the performance of *T. achaeae. Trichogramma achaeae* displayed a strong plastic response in fecundity to acclimation supporting our hypothesis that parasitoids would achieve the highest performance when acclimated at their optimal temperature. The highest fecundity was achieved for parasitoids acclimated at the optimal temperature of 25 °C and decreased sharply with decreasing acclimation temperature. This can been ascribed to an influence on egg maturation which is known to be greatly limited by low temperature (Steigenga and Fischer, 2007; Berger et al., 2008). Development time and body size contradicted the hypothesis that parasitoids would perform well at the optimal temperature, as both development rate and body size increased when acclimation temperature decreased to 15 °C or increased to 30 °C. However, albeit significant, the changes were relatively small and could, in addition, be partly questioned, as the results might have been affected by the small sample size at 15 °C. Unlike previous studies on *T. brassicae* (Lessard and Boivin, 2013), we did not find that acclimation at low temperatures increase the longevity of *T. achaeae* at low temperatures

Given parasitoids were from different generations and their quality may have altered during rearing and thus differed between experiments, direct comparison of results from the two experiments cannot be made. However, since the highest fecundity was almost equal in both experiments, it might be assumed that the reproductive capacity of *T. achaeae* was maintained among generations. Given this premise, and given the fact *T. achaeae* acclimated at 15 °C (Experiment 2) had a fecundity comparable to that of 23 °C-reared parasitoids tested at 15 °C (Experiment 1), our study has not supported the notion that cold acclimation will lead to an increased fecundity in cold environments as previously reported for *T. achaeae* by (Cascone et al., 2015). Since no significant reduction of performance was observed when parasitoids were acclimated at 30 °C, further studies regarding performance at extreme temperatures are needed to establish whether rearing at 30 °C will result in increased performance at a broad range of temperatures. This could improve rearing turnover and biological control efficacy in the field.

An ideal BCA should be able to maintain a relative high efficacy at a prolonged period under variable environments. This study found that *T. achaeae* will be able to survive and perform well at the temperature range encountered during most field applications. Considering the low fecundity at 15 °C, more parasitoids should be released to achieve the same efficacy compared to parasitoids applied at high temperatures. The possibility for altering performance through thermal acclimation might also have implication in relation to cold storage. Given cold acclimation reduced fecundity, it is not recommended to store *T. achaeae* in the form of parasitised host eggs at low temperature for a long time.

Quality control by periodically monitoring several parameters is recommended to maintain and improve quality of parasitoids and ensure field success. Seeing as fecundity was heavily weighted to the overall performance and was influenced by female proportion, measuring these two parameters are essential to quality control. Considering the differences between laboratory and field conditions, further studies linking laboratory assessment to field performance are required.

## Author contribution

**Long Chen**: methodology, investigation, data curation, formal analysis, writing-original draft. **Jesper Givskov Sørensen**: conceptualisation, methodology, validation, supervision writing-review & editing, funding acquisition. **Annie Enkegaard**: methodology, validation, supervision, writing-review & editing, funding acquisition.

## Acknowledgement

This work was supported by the Department of Agroecology, Aarhus University; and a grant from Aarhus University Research Foundation to JGS [AUFF-E-2015-FLS-8-72]. The authors would like to express special thanks to Dr. Xiangyu Guo, Centre for Quantitative Genetics and Genomics, Aarhus University, for her assistance on statistical analyses and to Dr. Kim Jensen, Department of Bioscience, Aarhus University, for helpful discussion of the manuscript.

